# Phylogeny, evidence for a cryptic plastid, and distribution of *Chytriodinium* parasites (Dinophyceae) infecting copepods

**DOI:** 10.1101/418467

**Authors:** Jürgen F. H. Strassert, Elisabeth Hehenberger, Javier del Campo, Noriko Okamoto, Martin Kolisko, Thomas A. Richards, Alexandra Z. Worden, Alyson E. Santoro, Patrick J. Keeling

## Abstract

Spores of the dinoflagellate *Chytriodinium* are known to infest copepod eggs causing their lethality. Despite the potential to control the population of such an ecologically important host, knowledge about *Chytriodinium* parasites is limited: we know little about phylogeny, parasitism, abundance, or geographical distribution. We carried out genome sequence surveys on four manually isolated sporocytes from the same sporangium to analyse the phylogenetic position of *Chytriodinium* based on SSU and concatenated SSU/LSU rRNA gene sequences, and also characterize two genes related to the plastidial heme pathway, *hemL* and *hemY*. The results suggest the presence of a cryptic plastid in *Chytriodinium* and a photosynthetic ancestral state of the parasitic *Chytriodinium*/*Dissodinium* clade. Finally, by mapping *Tara* Oceans V9 SSU amplicon data to the recovered SSU rRNA gene sequences from the sporocytes, we show that globally, *Chytriodinium* parasites are most abundant within the pico/nano- and mesoplankton of the surface ocean and almost absent within microplankton, a distribution indicating that they generally exist either as free-living spores or host-associated sporangia.

## INTRODUCTION

The marine zooplankton are dominated by copepods, which constitute a large source of animal protein globally, and are a major food source of numerous crustaceans, fish, and — beside krill — baleen whales. Both zooplankton fecal pellets and respiration at high depths sequester carbon to the deep sea, reducing the return of CO_2_ to the atmosphere (Steinberg et al. 2008; Jónasdóttir et al. 2015). Parasites of copepods are known to affect their population dynamics, but little is known about many of these parasites (Skovgaard 2014). Dinoflagellates of the genus *Chytriodinium* have been shown to be one such group of copepod parasites: their dinospores infest the lipid-rich eggs and, while building a cyst, which produces sporocytes that divide and form new flagellated spores, they absorb the eggs’ contents (e.g., Cachon and Cachon 1968; Gómez et al. 2009). Despite the potential to impact host populations, our knowledge of *Chytriodinium* is scarce. Morphological data are limited to light-microscopic investigations, and their apparently complex life cycle resulted in contradicting phylogenetic assignments at various taxonomic ranks within the Dinophyceae (see Gómez et al. 2009 and references within). Molecular sequence data should resolve this question, but sequence data for this genus is rare, and consequently, the most recently published SSU rRNA gene based tree illustrating its phylogenetic position contains only a single full-length and two partial *Chytriodinium* sequences (Gomez and Skovgaard 2015). This phylogeny suggests an affiliation of *Chytriodinium* to the *Gymnodinium* clade (Daugbjerg et al. 2000), and a split of this clade into free-living and parasitic subgroups; the latter comprising *Chytriodinium* and *Dissodinium* in the family Chytriodiniaceae. Monophyly of Chytriodiniaceae, however, was not statistically supported and could also not be confirmed using LSU rRNA gene sequences as phylogenetic marker (Gomez and Skovgaard 2015).

Members of the *Gymnodinium* clade show diverse trophic modes and plastids of various origins. Among the parasites, a plastid (containing chlorophyll *a*) has been reported in one of the two described *Dissodinium* species (*Dissodinium pseudolunula*; Gómez 2012). Nevertheless, based on the phylogeny, a recent loss of the plastids has been proposed for the Chytriodiniaceae and the presence of a plastid in *D. pseudolunula* interpreted as an indication of a photosynthetic ancestor and not as evidence for a recent plastid acquisition (Gómez 2012).

Although copepods are abundant and widespread in the ocean, studies on the geographic/habitat distribution and abundance of chitriodinids, which can give evidence for their capability to control the hosts’ occurrence, are not available. Furthermore, the absence of *Chytriodinium* in identification keys and the lack of highly distinctive features, such as the lunate sporangia in *D. pseudolunula*, makes it likely that this genus has not been recognized by researchers in earlier plankton surveys (Gomez and Skovgaard 2015).

In this study, we carried out genome sequencing from four parasites by manually isolating sporocytes matching the overall description of *Chytriodinium affine* as they were released from a cyst attached to a copepod nauplius collected from Monterey Bay. Genomic data allowed us to reassess the phylogeny of *Chytriodinium* by analysing the SSU and LSU rRNA gene sequences, collect evidence for the presence of a cryptic plastid, and by mapping *Tara* Oceans amplicon data to the SSU V9 region examine the abundance and ecological distribution of *Chytriodinium* species.

## MATERIAL AND METHODS

### Sampling and genomic data generation

Seawater was sampled from 60 m depth with Niskin bottles mounted on a rosette sampler in October 2014 in Monterey Bay (coordinates: 36°47′44.9″ N, 121°50′47.4″ W; for more details, see Strassert et al. 2018). Four sporocytes released from a copepod-attached cyst were manually isolated using micro-capillaries. Genomic DNA of each sporocyte was amplified with the REPLI-g UltraFast Mini Kit (Qiagen; protocol for blood cells, 16 h incubation) and the products were used for PCR-based SSU rRNA gene amplification using universal eukaryotic primers (Gile et al. 2011). SSU rRNA genes were cloned with the StrataClone PCR Cloning Kit (Agilent Technologies) and sequenced to identify the sporocytes’ phylogenetic affiliation. TruSeq library preparation and Illumina MiSeq sequencing (PE, 300 bp) were conducted at McGill University and Génome Québec Innovation Centre as described by Gawryluk et al. (2016). Reads have been submitted to GenBank under accession number: SRP129890. In addition to the four individually isolated sporocytes, five further sporocytes were collected and checked for their identity as described above (in total 17 clones), but not used for genome sequencing.

### Genome assembly, decontamination, and annotation

With exception of the genome assembly, the procedures were generally conducted as described elsewhere (Strassert et al. 2018). In short: the sequence data of the four samples (showing nearly identical SSU rRNA gene sequences) was pooled and FastQC (Andrews 2010) was used to evaluate the data quality. Reads were trimmed and merged using Trimmomatic (Bolger et al. 2014) and PEAR (Zhang et al. 2014), respectively, and quality-filtered with Sickle (Joshi and Fass 2011). The reads were assembled using Ray v2.3.1 (Boisvert et al. 2012) with a kmer of 67 and a minimum contig length of 150 bp. To identify and remove putative contaminations present in the assembly, the contigs were subjected to BLAST (Altschul et al. 1990) searches against the nt nucleotide database of NCBI as well as the Swiss-Prot database (Poux et al. 2016) of UniProt (E-value = 1e-25 for both searches). Additionally, the quality-filtered reads were mapped to the assembled contigs with bowtie2 v. 2.2.6 (Langmead and Salzberg 2012). The BLAST search results and the mapped reads were analyzed using blobtools v1.0 (Laetsch and Blaxter 2017) to identify contaminants (prokaryotic, viral and human sequences) and subsequently remove them together with contigs that had a read coverage of less than five reads (= clean dataset 1; 70.536 contigs). A second dataset was created by additionally removing all contigs that had no hit in either BLAST database and had a read coverage of less than five reads (= clean dataset 2; 3.268 contigs). These “no hit” contigs were of relatively short length (N50 = 664 nt) and were also defined by a low read coverage in general. Uncleaned and both cleaned datasets are available at the Dryad Digital Repository: ###. The assembly was checked for homologs in the KEGG database with KAAS (Moriya et al. 2007).

### Phylogenetic analyses

SSU rRNA gene sequences of the *Gymnodinium* clade and other selected taxa were exported from the SILVA123 database (Quast et al. 2013) and aligned together with the newly obtained sequence using MAFFT-L-INS-i v. 7.215 (Katoh and Standley 2013). The alignment was trimmed using trimAl v1.4 (Capella-Gutierrez et al. 2009) with the automated1 flag. One thousand maximum likelihood (ML) trees were reconstructed with RAxML v8.2.4 (Stamatakis 2014) using the GTRGAMMAI rate distribution model, and the tree topology was tested with 5,000 standard bootstrap replicates. A second tree was calculated as described above with exception that the concatenated SSU+LSU rRNA gene sequence alignment used fewer taxa due to the limited availability of LSU.

Two further maximum likelihood trees were inferred from protein sequences of the genes *hemL* and *hemY*. The candidates were used to query a custom protein database with BLAST (E-value threshold ≤1e-5). Initial parsing of the results was performed with an E-value of 1e-25 and a query coverage of 50%. Due to the low number of hits (nine), a relaxed E-value of 1e-5 was used for *hemY*. The parsed sequences were aligned with MAFFT-L-INS-i and trimmed with trimAl using a gap threshold of 20%. FastTree (Price et al. 2009) with default settings was used to reconstruct the initial phylogenies. The resulting trees and the underlying alignments were manually inspected to identify and remove contaminants from the alignments. Only the cyanobacterial clade of the initial *hemL* phylogeny was retained. The remaining sequences were re-aligned and only the domains of the respective proteins were used in the consecutive analyses; i.e., for *hemL*, the OAT_like conserved domain family cd00610 and for *hemY*, the Amino_oxidase Pfam family PF01593 (both as predicted for *Arabidopsis thaliana* orthologues present in the alignments). The domain alignments were trimmed as described above, resulting in a final length of 413 aa and 446 aa for *hemL* and *hemY*, respectively. Maximum likelihood trees were reconstructed with RAxML, calculating the best of 50 trees and 1,000 standard bootstrap replicates using the LG+Γ model.

All alignments used in this study are available on request.

### Abundance and distribution analyses

SSU rRNA V9 amplicons were recruited from the *Tara* Oceans OTU database (de Vargas et al. 2015) by BLASTN searches against the SSU rRNA gene sequence obtained in this study (sequence similarity cutoff: 99%). Geographical, size fraction (pico/nano: 0.8–20 μm, micro: 20–180 μm, and meso: 180–2,000 μm), depth, and temperature distributions were analysed using QIIME (Caporaso et al. 2010) and the metadata linked to the retrieved amplicons. Amplicon Numbers were normalized using CSS as describe elsewhere (http://qiime.org/).

## RESULTS AND DISCUSSION

We collected seawater from 60 m depth on Line 67 in the Monterey Bay, Northeastern Pacific Ocean, and microscopic examination of protist diversity revealed the presence of a copepod nauplius (identified by its extremities) with a large attached cyst (approximately 85 μm in diameter; Fig. 1A, B). Inside the cyst, sporocytes were packed in the form of a coiled chain built presumably as a result of palintomy (Cachon and Cachon 1968). Upon disruption, the chain of sporocytes (Fig. 1C) was released from the cyst, and individual sporocytes were separated from each other within five minutes. Four individual sporocytes were manually collected and morphologically documented under an inverted microscope while at sea and preserved for genomic DNA amplification. The sporocytes showed a division line in the middle (Fig. 1D) and their cytoplasm had fine hyaline granules, some of which were lightly pigmented in brownish-orange. The final products of the division, i.e., the flagellated dinospores, were not observed prior to isolation.

**Figure 1.**
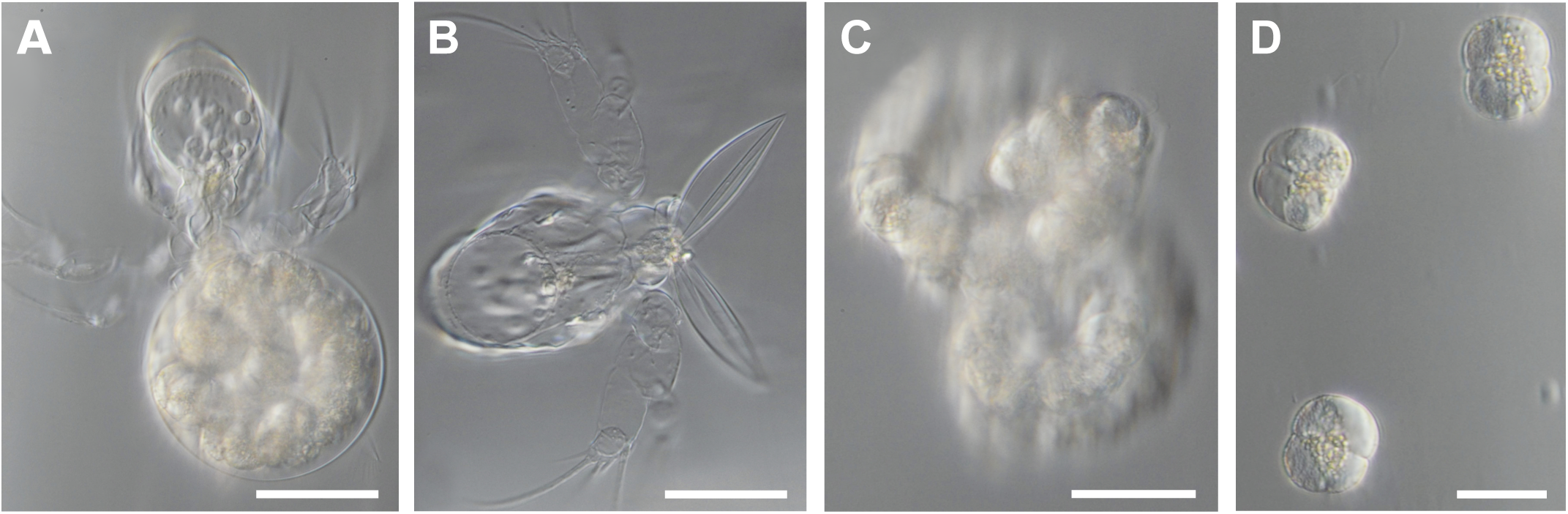
Morphology of *Chytriodinium affine*. The light micrographs show the parasite at different stages. A: Cyst containing a chain of sporocytes attached to a copepod. B: The same copepod (turned 90°) after cyst detachment. C: Coiled chain of sporocytes released from the ruptured cyst. D: Dividing sporocytes. Scale bars: A and B = 50 μm, C = 40 μm, D = 20 μm.

From the total DNA, we amplified the full-length SSU rRNA gene sequence, which resulted in a single sequence type from all four isolated sporocytes. In addition to the four sporocytes used in this study, the SSU rRNA genes of five further manually isolated sporocytes from the same cyst were sequenced and again all were nearly identical forming a clade (not shown) that is sister to a partial sequence (ca. 1,200 bp) previously characterized from *Chytriodinium affine* from the Mediterranean Sea (accession number FJ473380; Gómez et al. 2009). Our sequence type shared 99% identity with the Mediterranean Sea isolate. We therefore tentatively identify our isolate as *C. affine,* however, it is noteworthy that our isolate is a novel phylotype of *C. affine,* suggesting that distinct populations exist.

The SSU rRNA phylogenetic analysis revealed nine phylotypes that could be assigned to the genus *Chytriodinium* (Fig. 2). As expected, our isolates branched with *C. affine* (and one environmental sequence), while sequences from *Chytriodinium roseum* and several uncultured isolates obtained in other studies formed the remainder of the well-supported clade and branched with *Dissodinium* within the *Gymnodinium* clade (Daugbjerg et al. 2000). Monophyly of *Chytriodinium* and *Dissodinium* (Chytriodiniaceae) remained unsupported, but the same branching pattern has been seen consistently in both maximum likelihood and Bayesian analyses (Gómez et al. 2009; Gomez and Skovgaard 2015). A split into the parasitic Chytriodiniaceae and the free-living *Gymnodinium* clade members was not recovered, as also shown by Gómez et al. (2009) but conflicting with the tree inferred by Bayesian analysis (Gomez and Skovgaard 2015), so the question as to whether the *Gymnodinium* clade is bifurcated into parasites and free-living forms cannot be answered at present. The maximum likelihood tree inferred from concatenated SSU/LSU alignment of the *Gymnodinium* clade did not show such a split, but the nodes in question remained unsupported (Fig. S1). In addition, monophyly of Chytriodiniaceae could not be recovered in this tree, which may be explained by the low number of taxa and partial character of the sequences available for this group.

**Figure 2.**
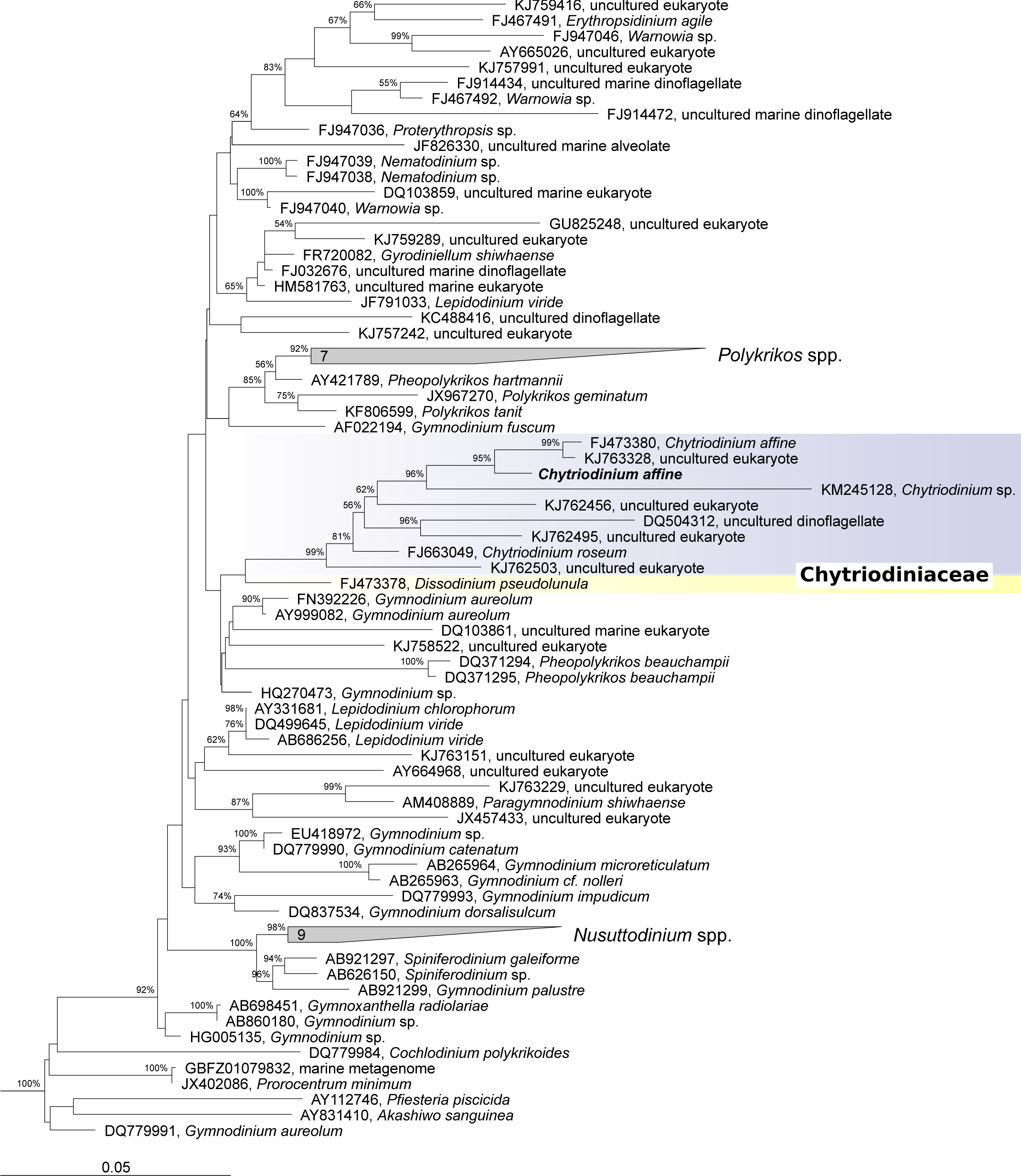
Phylogenetic position of *Chytriodinium affine*. The tree was inferred using maximum likelihood analysis of SSU rRNA gene sequences (>1,770 unambiguously aligned nucleotide positions; 4.4% gaps). Node support is shown by RAxML bootstraps (non-parametric). Numbers in polygons indicate the number of grouped taxa. The tree was rooted with representatives of the dinoflagellate genus *Alexandrium*.

We also examined the genome survey data for other genes and found identifiable genes encoding proteins relating to a wide variety of functions, as expected for a heterotrophic dinoflagellate (for a summary of KEGG hits, see Table S1). Specifically searching for genes related to plastid function revealed two likely candidates, *hemL* and *hemY* (both not encoded in the plastid genome). *hemL* encodes a glutamate-1-semialdehyde aminotransferase (GSA-AT). GSA-AT is used in the plastidial heme pathway and is present in all dinokaryotes (core dinoflagellates), even the non-photosynthetic ones such as *Dinophysis* and *Noctiluca* (Hehenberger et al. 2014; Janouškovec et al. 2017). Its sequence was almost complete but was lacking the N-terminus and thus any targeting information. However, a phylogenetic analysis revealed that this protein clearly clusters within a group of plastid-targeted *hemL* orthologues of other dinokaryotes corroborating the *Chytriodinium* origin (Fig. S2). In dinokaryotes, the synthesis of aminolevulinic acid is catalyzed by glutamyl tRNA reductase (GTR, encoded by *hemA*) and GSA-AT using glutamyl-tRNA as precursor. In contrast, in Perkinsozoa and early-branching dinoflagellates such as *Oxyrrhis* and *Hematodinium*, the synthesis is catalyzed by a single enzyme (aminolevulinic acid synthase, ALAS) and takes place in the mitochondrion and not in the plastid (Kořený et al. 2011; Danne et al. 2013; Gornik et al. 2015). The finding of GSA-AT suggests that *C. affine* possesses a cryptic plastid. In support of this, a further gene of the plastidial pathway was discovered, *hemY*, coding for a protoporphyrinogen oxidase (PPOX). The *hemY* also lacked its N-terminus, but in phylogenies it was affiliated with orthologues from other dinokaryotes (Fig. S3). The traces of plastids in *D. pseudolunula* (Gómez 2012) and now also *C. affine* are most consistent with a photosynthetic Chytriodiniaceae ancestor and the retention of a cryptic plastid with limited metabolic functions.

We also used these data to examine the distribution and abundance of *Chytriodinium* amplicons in the global ocean by mapping *Tara* Oceans V9 SSU rRNA gene data. Using a cutoff of 99% sequence similarity, the two most abundant phylotypes, *Chytriodinium affine* and *Chytriodinium* sp., accounted for ca. 13,750 and 126,200 amplicons, respectively. For perspective, the latter represents 0.2% of all amplicons that mapped to dinoflagellates. Both species were present in pico/nano- and mesoplankton but almost absent in microplankton size fractions (Fig. 3A, B), and the highest abundance could be observed within the oceanic mixed layer at depths between 118 m and 148 m (despite sampling representation bias; see Fig. 3C). The size distribution is in agreement with the different prevalent life stages of *Chytriodinium*, i.e., free-living spores and host-associated sporangia. In this context, it is noteworthy that in this study, the sporangium seemed to be attached to a copepod nauplius and not to an egg or egg sac of a brood-carrying copepod species (Gomez and Skovgaard 2015). Unfortunately, however, due to difficult conditions at sea, the authors failed to take more high quality pictures and to further investigate the characteristics of a feeding tube connecting host and parasite. Thus, whether *Chytriodinium* infects not only copepod eggs but also larval stages, cannot be finally answered and will have to be confirmed or rejected in other studies. The increased occurrence of *Chytriodinium* in deep chlorophyll maximum (DCM) and even more in the mixed layer may reflect the distribution of the host copepods, as it is known that several species can be most frequently found at these depths (Longhurst 2007).

**Figure 3.**
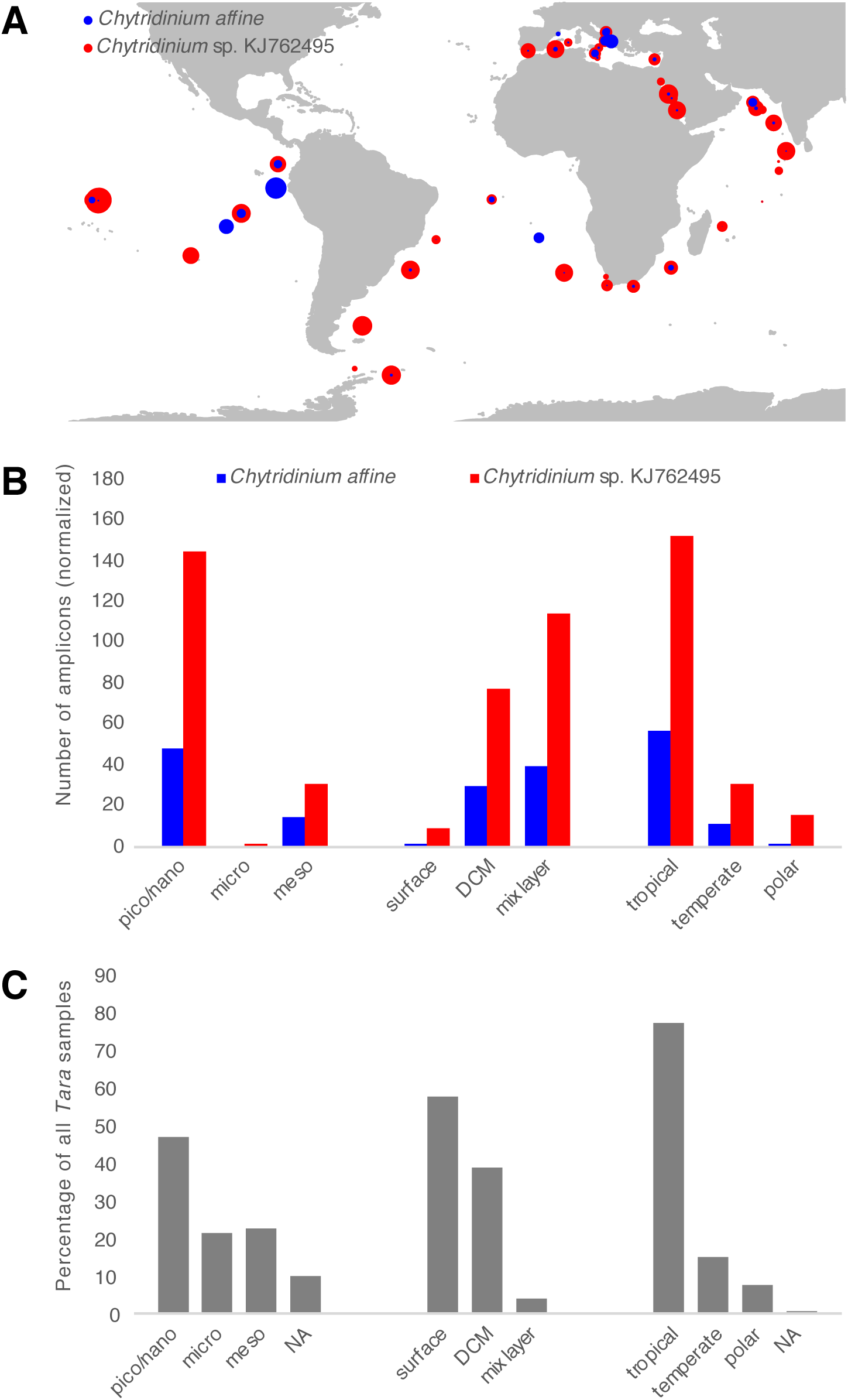
Distribution of *Chytriodinium affine* and *Chytriodinium* sp. (KJ762495) in *Tara* Oceans samples where they were detected in SSU V9 data using a sequence similarity cutoff of 99%. A: Geographical distribution. Dot sizes are proportional to the sum of the total amplicons at each location for the two species (detected in 169 of 335 *Tara* samples). Note that North Pacific data from *Tara* are not available. B: Size fraction, depth and temperature distributions. The abundances are based on normalized numbers of *Tara* Oceans V9 amplicons (the numbers reflect averages of samples where the two species were detected). C: Percentages of all *Tara* samples obtained from different size fractions, depths and water temperatures. N/A – information on size or temperature was not available; polar: <10 °C, temperate: 10–19 °C, tropical: >19 °C.

The patterns observed here for the global distribution of *Chytriodinium* parasites suggest they may play an important role in copepod population dynamics, and by extension impact marine food webs. Further investigations, in particular those focusing on their host interactions, will be of interest to fully uncover the ecological importance of these parasites.

## ACKNOWLEDGMENTS

We thank the captain and crew of the R/V *Western Flyer*, M. Blum, F. Chavez, V. Jimenez, J. T. Pennington, J. M. Smith, S. Sudek, J. Swalwell, C. Wahl, and S. Wilken, for logistical assistance prior to and during the cruises. We thank T. Glatzel and H.-D. Franke for identifying the host animal. This work was supported by a grant from the Gordon and Betty Moore Foundation (GBMF3307) to P.J.K., T.A.R., A.Z.W., and A.E.S. Ship time was supported by a grant from the David and Lucile Packard Foundation through MBARI and GBMF3788 to A.Z.W. J.dC. was supported by a Marie Curie International Outgoing Fellowship grant (FP7-PEOPLE-2012-IOF – 331450 CAARL), and N.O., M.K., and J.dC. were supported by a grant from the Tula Foundation to the Centre for Microbial Biodiversity and Evolution at UBC.

## SUPPORTING INFORMATION

**Table S1** Function of *Chytriodinium* genes based on KEGG annotation. KEGG Orthology (KO) numbers and their respective protein names are shown. Sheet 2 shows the results for “clean dataset 2” (see Methods).

**Figure S1** Phylogenetic tree inferred from concatenated SSU and LSU rRNA gene sequences of the *Gymnodinium* clade. The tree was reconstructed using maximum likelihood analysis of >3,540 unambiguously aligned nucleotide positions (13.4% gaps). Node support is shown by RAxML bootstraps (non-parametric). Sequences of *Alexandrium* were used as outgroup (not shown).

**Figure S2** Phylogenetic tree inferred from the maximum likelihood analysis of the OAT_like conserved domain (cd00610) of GSA-AT (glutamate-1-semialdehyde aminotransferase; encoded by *hemL*) of selected representative taxa. Black dots correspond to >95% ML bootstrap support. Numbers at nodes represent bootstrap supports of >50 %. Sequences were obtained from GenBank and MMETSP. For more details, see main text.

**Figure S3** Maximum likelihood tree based on the analysis of the Amino_oxidase domain (PF01593) of protoporphyrinogen oxidase (encoded by *hemY*) of diverse representatives. Black dots correspond to >95% ML bootstrap support. Numbers at nodes represent bootstrap supports of >50%. Sequences were obtained from GenBank and MMETSP. For more details, see main text.

## LITERATURE CITED

Altschul S. F., Gish W., Miller W., Myers E. W. & Lipman D. J. 1990. Basic local alignment search tool. J Mol Biol, 215:403–410.

Andrews S. 2010. FastQC: a quality control tool for high throughput sequence data. http://www.bioinformatics.babraham.ac.uk/projects/fastqc.

Boisvert S., Raymond F., Godzaridis É., Laviolette F. & Corbeil J. 2012. Ray Meta: scalable de novo metagenome assembly and profiling. Genome Biol, 13:R122.

Bolger A. M., Lohse M. & Usadel B. 2014. Trimmomatic: a flexible trimmer for Illumina sequence data. Bioinformatics, 30:2114–2120.

Cachon J. & Cachon M. 1968. Cytologie et cycle évolutif des *Chytriodinium*. Protistologica, 4:249–262.

Capella-Gutierrez S., Silla-Martinez J. M. & Gabaldon T. 2009. trimAl: a tool for automated alignment trimming in large-scale phylogenetic analyses. Bioinformatics, 25:1972–1973.

Caporaso J. G., Kuczynski J., Stombaugh J., Bittinger K., Bushman F. D., Costello E. K., Fierer N., Pena A. G., Goodrich J. K., Gordon J. I., Huttley G. A., Kelley S. T., Knights D., Koenig J. E., Ley R. E., Lozupone C. A., McDonald D., Muegge B. D., Pirrung M., Reeder J., Sevinsky J. R., Turnbaugh P. J., Walters W. A., Widmann J., Yatsunenko T., Zaneveld J. & Knight R. 2010. QIIME allows analysis of high-throughput community sequencing data. Nat Methods, 7:335–336.

Danne J. C., Gornik S. G., MacRae J. I., McConville M. J. & Waller R. F. 2013. Alveolate mitochondrial metabolic evolution: dinoflagellates force reassessment of the role of parasitism as a driver of change in apicomplexans. Mol Biol Evol, 30:123–139.

Daugbjerg N., Hansen G., Larsen J. & Moestrup Ø. 2000. Phylogeny of some of the major genera of dinoflagellates based on ultrastructure and partial LSU rDNA sequence data, including the erection of three new genera of unarmoured dinoflagellates. Phycologia, 39:302–317.

Gawryluk R. M. R., Del Campo J., Okamoto N., Strassert J. F. H., Lukes J., Richards T. A., Worden A. Z., Santoro A. E. & Keeling P. J. 2016. Morphological identification and single-cell genomics of marine diplonemids. Curr Biol, 26:3053–3059.

Gile G. H., James E. R., Scheffrahn R. H., Carpenter K. J., Harper J. T. & Keeling P. J. 2011. Molecular and morphological analysis of the family Calonymphidae with a description of *Calonympha chia* sp. nov., *Snyderella kirbyi* sp. nov., *Snyderella swezyae* sp. nov. and *Snyderella yamini* sp. nov. Int J Syst Evol Microbiol, 61:2547–2558.

Gómez F. 2012. A quantitative review of the lifestyle, habitat and trophic diversity of dinoflagellates (Dinoflagellata, Alveolata). Syst Biodivers, 10:267–275.

Gómez F., Moreira D. & López-García P. 2009. Life cycle and molecular phylogeny of the dinoflagellates *Chytriodinium* and *Dissodinium*, ectoparasites of copepod eggs. Eur J Protistol, 45:260–270.

Gomez F. & Skovgaard A. 2015. Molecular phylogeny of the parasitic dinoflagellate *Chytriodinium* within the *Gymnodinium* Clade (Gymnodiniales, Dinophyceae). J Eukaryot Microbiol, 62:422–425.

Gornik S. G., Febrimarsa, Cassin A. M., MacRae J. I., Ramaprasad A., Rchiad Z., McConville M. J., Bacic A., McFadden G. I., Pain A. & Waller R. F. 2015. Endosymbiosis undone by stepwise elimination of the plastid in a parasitic dinoflagellate. Proc Natl Acad Sci U S A, 112:5767–5772.

Hehenberger E., Imanian B., Burki F. & Keeling P. J. 2014. Evidence for the retention of two evolutionary distinct plastids in dinoflagellates with diatom endosymbionts. Genome Biol Evol, 6:2321–2334.

Janouškovec J., Gavelis G. S., Burki F., Dinh D., Bachvaroff T. R., Gornik S. G., Bright K. J., Imanian B., Strom S. L., Delwiche C. F., Waller R. F., Fensome R. A., Leander B. S., Rohwer F. L. & Saldarriaga J. F. 2017. Major transitions in dinoflagellate evolution unveiled by phylotranscriptomics. Proc Natl Acad Sci U S A, 114:E171–E180.

Jónasdóttir S. H., Visser A. W., Richardson K. & Heath M. R. 2015. Seasonal copepod lipid pump promotes carbon sequestration in the deep North Atlantic. Proc Natl Acad Sci U S A, 112:12122–12126.

Joshi N. A. & Fass J. N. 2011. Sickle: a sliding-window, adaptive, quality-based trimming tool for FastQ files (Version 1.33) [Software]. https://github.com/najoshi/sickle.

Katoh K. & Standley D. M. 2013. MAFFT multiple sequence alignment software version 7: improvements in performance and usability. Mol Biol Evol, 30:772–780.

Kořený L., Sobotka R., Janouškovec J., Keeling P. J. & Oborník M. 2011. Tetrapyrrole synthesis of photosynthetic chromerids is likely homologous to the unusual pathway of apicomplexan parasites. Plant Cell, 23:3454–3462.

Laetsch D. R. & Blaxter M. L. 2017. BlobTools: interrogation of genome assemblies. F1000Research, 6:1287.

Langmead B. & Salzberg S. L. 2012. Fast gapped-read alignment with Bowtie 2. Nat Methods, 9:357–359.

Longhurst A. R. 2007. Fronts and pycnoclines: ecological discontinuities. *In*: Longhurst A. R. (ed.), Ecological Geography of the Sea. Burlington, Academic Press. p. 35–49.

Moriya Y., Itoh M., Okuda S., Yoshizawa A. C. & Kanehisa M. 2007. KAAS: an automatic genome annotation and pathway reconstruction server. Nucleic Acids Res, 35:W182–5.

Poux S., Arighi C. N., Magrane M., Bateman A., Wei C.-H., Lu Z., Boutet E., Bye-A-Jee H., Famiglietti M. L. & Roechert B. 2016. On expert curation and sustainability: UniProtKB/Swiss-Prot as a case study. bioRxiv.

Price M. N., Dehal P. S. & Arkin A. P. 2009. FastTree: computing large minimum evolution trees with profiles instead of a distance matrix. Mol Biol Evol, 26:1641–1650.

Quast C., Pruesse E., Yilmaz P., Gerken J., Schweer T., Yarza P., Peplies J. & Glockner F. O. 2013. The SILVA ribosomal RNA gene database project: improved data processing and web-based tools. Nucleic Acids Res, 41:D590–6.

Skovgaard A. 2014. Dirty tricks in the plankton: diversity and role of marine parasitic protists. Acta Protozool, 53:51–62.

Stamatakis A. 2014. RAxML version 8: a tool for phylogenetic analysis and post-analysis of large phylogenies. Bioinformatics, 30:1312–1313.

Steinberg D. K., Van Mooy B. A. S., Buesseler K. O., Boyd P. W., Kobari T. & Karl D. M. 2008. Bacterial vs. zooplankton control of sinking particle flux in the ocean’s twilight zone. Limnol Oceanogr, 53:1327–1338.

Strassert J. F. H., Karnkowska A., Hehenberger E., Del Campo J., Kolisko M., Okamoto N., Burki F., Janouskovec J., Poirier C., Leonard G., Hallam S. J., Richards T. A., Worden A. Z., Santoro A. E. & Keeling P. J. 2018. Single cell genomics of uncultured marine alveolates shows paraphyly of basal dinoflagellates. ISME J, 12:304–308.

de Vargas C., Audic S., Henry N., Decelle J., Mahe F., Logares R., Lara E., Berney C., Le Bescot N., Probert I., Carmichael M., Poulain J., Romac S., Colin S., Aury J.-M., Bittner L., Chaffron S., Dunthorn M., Engelen S., Flegontova O., Guidi L., Horak A., Jaillon O., Lima-Mendez G., Lukes J., Malviya S., Morard R., Mulot M., Scalco E., Siano R., Vincent F., Zingone A., Dimier C., Picheral M., Searson S., Kandels-Lewis S., Acinas S. G., Bork P., Bowler C., Gorsky G., Grimsley N., Hingamp P., Iudicone D., Not F., Ogata H., Pesant S., Raes J., Sieracki M. E., Speich S., Stemmann L., Sunagawa S., Weissenbach J., Wincker P. & Karsenti E. 2015. Ocean plankton. Eukaryotic plankton diversity in the sunlit ocean. Science, 348:1261605.

Zhang J., Kobert K., Flouri T. & Stamatakis A. 2014. PEAR: a fast and accurate Illumina Paired-End reAd mergeR. Bioinformatics, 30:614–620.

